# Machine Learning of the Cardiac Phenome and Skin Transcriptome to Categorize Heart Disease in Systemic Sclerosis

**DOI:** 10.1101/213678

**Authors:** Monique E. Hinchcliff, Tracy M. Frech, Tammara A. Wood, Chiang-Ching Huang, Jungwha Lee, Kathleen Aren, John J. Ryan, Brent Wilson, Lauren Beussink-Nelson, Michael L. Whitfield, Rahul C. Deo, Sanjiv J. Shah

**Affiliations:** From the Divisions of Rheumatology University of California—San Francisco, San Francisco, CA; From the Divisions of Rheumatology and Cardiology University of California—San Francisco, San Francisco, CA; Department of Medicine, and Department of Preventive Medicine University of California—San Francisco, San Francisco, CA; Northwestern University Feinberg School of Medicine, Chicago, IL; Division of Rheumatology University of California—San Francisco, San Francisco, CA; Northwestern University Feinberg School of Medicine, Chicago, IL; Division of Rheumatology and Cardiology University of California—San Francisco, San Francisco, CA; Department of Medicine, University of Utah, Salt Lake City, UT; Department of Molecular and Systems Biology, Geisel School of Medicine at Dartmouth, Hannover, NH University of California—San Francisco, San Francisco, CA; Zilber School of Public Health, University of Wisconsin-Milwaukee, Milwaukee, WI University of California—San Francisco, San Francisco, CA; Division of Cardiology, Department of Medicine, Cardiovascular Research Institute, and California Institute for Quantitative Biosciences, University of California—San Francisco, San Francisco, CA

**Author notes:** Address for correspondence: Sanjiv J. Shah, MD Division of Cardiology Department of Medicine Feinberg Cardiovascular Research Institute Northwestern University Rahul C. Deo, MD, PhD Division of Cardiology Department of Medicine Cardiovascular Research Institute University of California, San Francisco. **Sources of funding:** This work was supported by National Institutes of Health grants K23 AR059763 and L30 AR054311 (to M.E.H.), R01 HL107557 and R01 HL127028 (to S.J.S.), DP2 HL123228 (to R.C.D), and P60 AR064464 (to J.L.); American Heart Association grants #16SFRN28780016 and 15CVGPSD27260148 (to S.J.S.); and Scleroderma Research Foundation grants (to M.E.H and M.L.W.). **Disclosures:** None of the authors have any relationships with industry that are relevant to this manuscript.

**Keywords:** systemic sclerosis, echocardiography, skin, machine learning, gene expression

## Abstract

**Background:** Cardiac involvement is a leading cause of death in systemic sclerosis (SSc/scleroderma). The complexity of SSc cardiac manifestations is not fully captured by the current clinical SSc classification, which is based on extent of skin involvement and specific autoantibodies. Therefore, we sought to develop a clinically relevant SSc cardiac disease classification to improve clinical care and increase understanding of SSc cardiac disease pathobiology. We hypothesized that machine learning could identify novel SSc cardiac disease subgroups, and that gene expression assessment of skin could provide insights into molecular pathogenesis of these SSc pheno-groups.

**Methods:** We used unsupervised model-based clustering (phenomapping) of SSc patient echocardiographic and clinical data to identify clinically relevant SSc pheno-groups in a discovery cohort (n=316), and validated these findings in an external SSc validation cohort (n=67). Cox regression was used to evaluate survival differences among groups. Gene expression profiles from skin biopsies from a subset of SSc patients (n=68) and controls (n=18) were analyzed with weighted gene co-expression network analyses to identify gene modules that were associated with cardiac pheno-groups and echocardiographic parameters.

**Results:** Four SSc cardiac pheno-groups were identified with distinct profiles. Pheno-group #1 displayed a predominant cutaneous phenotype without cardiac involvement; pheno-group #2 had long-standing SSc with limited skin and cardiac involvement; pheno-group #3 had diffuse skin involvement, a high frequency of interstitial lung disease (88%), and significant right heart remodeling/dysfunction; and pheno-group #4 had prolonged SSc disease duration, limited skin involvement, and marked biventricular cardiac involvement. After multivariable adjustment, pheno-group #3 (hazard ratio [HR] 7.8, 95% confidence interval [CI] 1.5–33.0) and pheno-group #4 (HR 10.5, 95% CI 2.1–52.7) remained associated with mortality (P<0.05). The addition of pheno-group classification was additive to conventional survival models (P<0.05 by likelihood ratio test for all models), a finding that was replicated in the validation cohort. Skin gene expression analysis identified 2 gene modules (representing fibrosis and skin integrity, respectively) that differed among the cardiac pheno-groups and were associated with specific echocardiographic parameters.

**Conclusions:** Machine learning of echocardiographic and skin gene expression data in SSc identifies clinically relevant subgroups with distinct cardiac phenotypes, survival, and associated molecular pathways in skin.

## INTRODUCTION

Systemic sclerosis (SSc)/scleroderma—an autoimmune disease characterized by inflammation and autoantibody production, vascular perturbations, and fibrosis—is a compelling example of a poorly understood, heterogeneous, systemic disease that is in need of novel effective treatments.^1^ Cardiac involvement in SSc is common, but its manifestations are highly variable and diverse.^2, 3^ Various classification systems for SSc have been developed to help resolve SSc disease heterogeneity. Current methods include patient stratification into limited cutaneous (lcSSc) and diffuse cutaneous (dcSSc) subsets based on degree of skin fibrosis (i.e., modified Rodnan skin score [mRSS]);^4^ classification based on serum autoantibodies;^5^ and an intrinsic subset classification, which distinguishes 4 molecular SSc subgroups (inflammatory, limited, normal-like, and fibroproliferative) based upon gene expression in skin biopsies.^6^ The mRSS and serum autoantibody classifications are helpful in broadly assessing risk for SSc internal organ manifestations and disease course while classification according to skin gene expression may be useful in identifying patients who are most likely to respond to current immune modulatory and anti-fibrotic therapies.^7–11^ Despite their usefulness, no SSc classifier reliably identifies SSc patients with specific types of cardiac involvement. Moreover, our understanding of SSc cardiac disease heterogeneity and pathogenesis is incomplete.^2, 3^

Cardiovascular manifestations in SSc include right and left ventricular systolic and diastolic dysfunction, pulmonary arterial and venous hypertension, pericardial disease, and life-threatening arrhythmias.^2, 3, 12–14^ Patients with SSc routinely undergo annual echocardiography with Doppler and tissue Doppler imaging to screen for and monitor progression of cardiopulmonary disease.^15–17^ This wealth of data is available and can be used to define discrete phenotypic groups (pheno-groups).^18^ Such a classification system in SSc would be clinically useful because long term SSc outcomes and mortality risk secondary to SSc heart disease could be more accurately estimated, failed clinical trials could be reanalyzed to determine if a subset of participants responded according to pheno-group, and insights could be obtained regarding the pathobiology of SSc cardiac disease.

Unsupervised learning is a category of machine learning that identifies inherent patterns within datasets.^19, 20^ Such approaches have been widely applied to genomic data to classify patients with cancer and autoimmune diseases.^21, 22^ However, for many organs (including the heart), genomic data is neither readily attainable nor likely to be sufficient to characterize the higher-order structural or functional changes that can strongly predict disease course and outcomes.^19, 23^ We have previously applied unsupervised clustering approaches to dense echocardiographic data for over 500 patients with heart failure and preserved ejection fraction, another challenging heterogeneous disease without known effective disease-modifying therapies.^18^ We identified discrete phenotypic clusters that differed substantially in clinical characteristics and outcomes, findings which suggest that unsupervised learning of phenotypic data (i.e., phenomapping) can be successfully applied to improve classification of heterogeneous disorders.

We therefore sought to apply an unsupervised learning approach to dense echocardiographic data in patients with SSc in order to develop a novel and informative cardiac disease classification scheme. Furthermore, because skin fibrosis is the predominant SSc clinical manifestation, we conducted parallel machine learning analyses of skin genome-wide gene expression data to investigate the molecular pathways underlying the differences among the identified cardiac pheno-groups. We hypothesized that distinct cardiac pheno-groups exist and differ substantially in clinical characteristics and survival, and are associated with distinct deregulated molecular pathways in skin.

## METHODS

### Study design

**Figure 1** displays the overall study design, which included unsupervised machine learning (model-based clustering) of echocardiographic and clinical phenotypes (i.e., phenomapping) in a training (discovery) cohort with testing in an external validation cohort. Clinical and echocardiographic characteristics, and survival, were subsequently compared among the resultant cardiac pheno-groups. In parallel, we conducted gene expression analyses of arm and back skin biopsies from SSc patients and healthy control subjects. Transcripts (probe sets) with large variability across control samples were excluded in order to restrict analyses to SSc-specific genes. Finally, the SSc skin gene modules were compared to SSc cardiac pheno-groups to determine which gene modules were most associated with the pheno-groups. For the significant SSc skin gene modules, we determined the clinical characteristics and echocardiographic parameters that were most associated with the identified gene modules.

**Figure 1.**
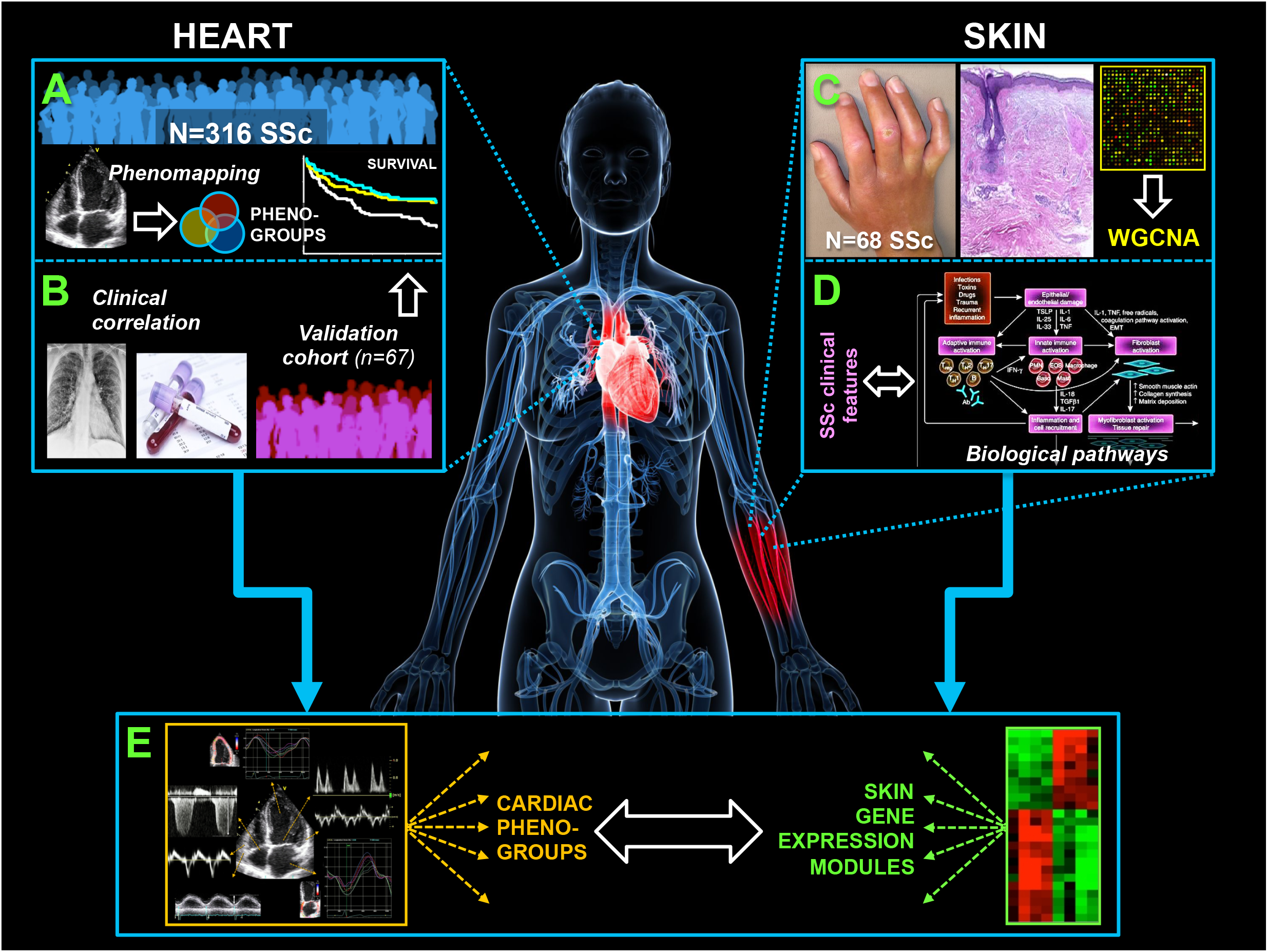
Machine Learning of the Cardiac Phenome and Skin Transcriptome to Resolve the Heterogeneity of Systemic Sclerosis-Associated Cardiac Involvement. Our research design included integrated analyses of cardiac echocardiographic and skin genomic data combined with conventional clinical phenotypic and disease outcome measures. (A) Unsupervised learning applied to quantitative echocardiographic data was used to identify coherent clusters of SSc patients with distinct cardiac disease phenotypes (echo-pheno-groups), and mortality was compared among the identified pheno-groups. (B) Next, we analyzed the clinical differences among the identified pheno-groups and replicated our classification system (and its association with survival) in an external validation cohort. (C) We subsequently analyzed gene expression from clinically affected and non-clinically affected skin from a subset of SSc patients, and also from healthy controls and used weighted-gene co-expression network analyses (WGCNA) to identify relevant gene modules. (D) Gene modules were then associated with SSc clinical features. (E) Finally, the gene modules were associated with the cardiac pheno-groups and also specific echocardiography phenotypes to integrate the phenomic and transcriptomic analyses.

### Discovery cohort

Between March 2007 and May 2011, 316 consecutive patients who fulfilled SSc diagnostic criteria^24^ underwent 2-dimensional echocardiography with Doppler and tissue Doppler imaging according to a standardized protocol based on published guidelines.^25, 26^ All patients were prospectively enrolled into the Northwestern Scleroderma Patient Registry and Biorepository, and all patients gave written, informed consent. The institutional review board at Northwestern University approved the study.

### Echocardiographic data

Echocardiograms conducted at the time of the initial clinical visit at the Northwestern Scleroderma Program were examined. Details of the echocardiographic protocol are provided in the **Supplementary Material** section. An experienced research sonographer (LBN) performed quantitative echocardiographic measurements, and a cardiologist with expertise in echocardiography (SJS) confirmed measurements. **Supplementary Table S1** lists the 56 continuous echocardiographic variables available for inclusion in machine learning analyses for the definition of SSc cardiac pheno-groups.

### Clinical variables

Clinical data within 3 months of the initial clinical evaluation, including age, height, weight, blood pressure, serum autoantibodies, mRSS (a semi-quantitative marker of SSc skin disease severity), SSc disease subtype, and SSc disease duration (defined as duration between first non-Raynaud SSc symptom and echocardiogram date) were collected by querying the Northwestern Enterprise Data Warehouse (comprehensive electronic health record) or by manual chart review.^27^

Patient-reported race was collected using a standardized data collection form. Diagnostic (pulmonary function tests [PFTs], high-resolution computed tomography of the chest [HRCT], cardiac catheterization, and chest radiography) exams were used to classify patients as having interstitial lung disease (ILD) and/or pulmonary artery hypertension (PAH) based on published guidelines.^28, 29^ Follow-up duration was defined as the duration between the echocardiogram and the last patient encounter or date of death within the electronic health record or the Social Security Death Index.

### Model-based clustering using a combination of continuous and binary variables

We sought an unsupervised learning approach that could: (1) accommodate both continuous and discrete variables; and (2) allow a principled method to select the number of distinct patient groups with similar cardiac disease (clusters). We selected the model-based approach implemented in the *flexmix* R package,^30^ which allows clusters to be defined by both continuous variables, as defined by a Gaussian distribution, and binary variables, as defined by a binomial distribution. The expectation-maximization algorithm was used to fit all parameters. The Bayesian information criterion, a penalty for model complexity, was applied to select the optimal number of clusters.

Missing echocardiography values were imputed with the SVDimpute function within the *impute* package in R. Next, to account for redundant echocardiographic variables, we performed principal components analysis (PCA), a form of linear dimensionality reduction. Principal component analysis uses linear combinations of the original variables to obtain a series of orthogonal (i.e. independent) features that are ranked in terms of the proportion of variability across the dataset they can explain. The top 6 principal components, collecting accounting for 65% of the variance, were used for clustering. We also noted a high correlation among several PFT variables; thus, PFT variables that were correlated at a correlation coefficient of >0.8 were filtered (keeping the PFT variable that was most informative and had the least missingness).

In addition to these cardiopulmonary indices of structure and function, we added 6 categorical variables: sex, race (white/non-white), mRSS (ordinal for each of 17 body areas, resulting in an integer score [0–51]), and positivity for 3 SSc-specific serum autoantibodies: anti-topoisomerase I (Scl-70), anti-RNA polymerase III, and anticentromere antibody because these variables have clinical significance in SSc.^5^ **Supplementary Table S1** lists all variables used for the machine learning analyses. Internal validation of the pheno-groups was conducted by re-running the *flexmix* expectation-maximization (EM) algorithm 50 times, and determining the most commonly predicted number of pheno-groups.

### Comparison of clinical characteristics and survival among pheno-groups

Once cluster membership (i.e., pheno-group) was defined, we compared differences in demographic, clinical, laboratory, PFT, and echocardiographic characteristics among the clusters (pheno-groups). Chi-squared tests (or Fisher exact tests, when appropriate) were used for categorical variables, and one-way analysis of variance (or Kruskal-Wallis test, when appropriate) was used for continuous variables.

To determine the clinical relevance of our phenomapping classification scheme, we used unadjusted and multivariable-adjusted Cox proportional hazards models to determine the independent association between pheno-groups and survival. The following covariates, which were selected based on clinical relevance, were included in a series of adjusted models: Model 1 included age, sex, SSc subtype (lcSSc vs. dcSSc), and SSc disease duration; Model 2 included all Model 1 covariates plus race, ILD, and PAH; and Model 3 included all Model 2 covariates plus the top 6 echocardiographic principal components. The proportionality assumption was tested and verified for all Cox regression models. Statistical analyses for the association of pheno-groups with outcomes were performed using Stata v.12.0 (StataCorp, College Station, TX).

We sought to further demonstrate the clinical importance of the SSc cardiac pheno-group classification by using alternate survival models to supplement our results from Cox regression analysis. We hypothesized that supervised learning methods that permit more complex variable combinations may be more effective at modeling survival than Cox proportional hazards models, which are restricted to a linear combination of predictors and as well as the proportional hazards assumption. This may be especially important in SSc since there are complex combinations of features that underlie high-risk SSc phenotypic groups (cardiac phenotypes, pulmonary phenotypes, serum autoantibody profiles, etc.). Therefore, we evaluated random survival forests (a supervised learning technique), which is an adaption of the random forests algorithm for right-censored data^31^ and compared the results to our Cox regression models (see **Supplementary Material** for additional details regarding the methods used for the random survival forests model).

### Validation cohort

After permission was obtained from the University of Utah Institutional Review Board, de-identified echocardiograms and clinical data were obtained for 67 subjects that fulfilled SSc criteria and had an echocardiogram obtained at the University of Utah. These subjects were prospectively enrolled and followed for outcomes at the University of Utah Scleroderma Program between January 2012 and February 2014. The same investigator (LBN) that scored Northwestern Scleroderma Program echocardiograms quantified the Utah echocardiograms, and SJS confirmed measurements. Demographic, clinical, and echocardiographic characteristics were compared between the discovery (Northwestern) and validation (Utah) cohorts. Student t-tests (or Wilcoxon rank-sum tests, when appropriate) were used for comparisons of continuous variables, and chi-squared tests were used for comparisons of categorical variables between the 2 groups.

Given that the validation cohort lacked some of the phenotypic measures that were available for our discovery cohort (specifically PFT data), we developed a regression model (using lasso)^32^ for discrimination of low-risk (pheno-groups #1 and #2) and high-risk (pheno-groups #3 and #4) SSc clusters using the remaining variables, and applied this to the validation cohort.

### Microarray processing of skin biopsies and analysis

Skin gene expression data were obtained from the subset of SSc patients at Northwestern University that had undergone skin biopsies and echocardiograms in order to identify biological processes that may underlie the identified pheno-groups. Skin biopsies from the clinically involved dorsal surface of the forearm and the clinically uninvolved flank/back were obtained from SSc subjects as well as from healthy control subjects, and microarray studies were performed as previously described,^33^ and as described in detail in the **Supplementary Material** section.

### Selection of SSc-variable transcripts

Skin biopsy samples from healthy control subjects recruited at Northwestern University were used to identify the most highly variable transcripts between forearm and flank/back biopsies unrelated to the SSc disease process.^11^ Coefficients of variation were computed for each transcript by dividing the mean intensity by the standard deviation across all control samples. A 75th percentile threshold was used to define a “non-disease variable” set (see **Supplementary Material** section).

### Weighted gene co-expression network analysis (WGCNA)

We focused on the 23,530 probe sets shared between the SSc samples and control samples. We eliminated any overlap with the non-disease variable set and identified the 2,500 (top 10^th^ percentile) most variable transcripts (referred as the “SSc variable” set), as judged by the coefficient of variation (see **Supplementary Material**).

To find biologically coherent gene expression modules within the SSc discovery cohort, we employed Weighted Gene Correlation Network Analysis (WGCNA). This commonly used dimensionality reduction method applies hierarchical clustering to a transformed correlation matrix of gene expression profiles in order to define highly correlated/covarying gene sets.^22, 34^ WGCNA was performed on the SSc variable set using the WGCNA package in R.^34^ Module activity for each sample was computed using the factor loading of the first principal component of the matrix of expression values for genes in the module (described as the first “eigengene” in WGCNA).

### Gene ontology enrichment analysis

Gene ontology (GO) enrichment was performed in order to identify the molecular pathways to which differentially expressed genes/genes modules belonged. Funcassociate 2.1 was used,^35^ which uses a Fisher’s exact test for enrichment, and permutation based correction for multiple hypothesis testing. Gene lists were filtered to only include genes with one or more mapped GO terms.

### Association of gene expression module activity with SSc clinical characteristics, echocardiographic phenotypes, and pheno-groups

The association of gene module activities and clinical and echocardiographic traits, and pheno-groups, was examined using linear regression and logistic regression models. Quantitative predictor variables were divided by twice the standard deviation to allow comparison of coefficients between continuous and binomial predictors.^36^

## RESULTS

### Model-based clustering identifies 4 novel SSc cardiac pheno-groups

The discovery cohort consisted of n=316 consecutive SSc patients, and the external validation cohort consisted of n=67 consecutive SSc patients. As shown in **Table 1**, in the discovery cohort, the mean age was 51±12 years, 84% were female, 29% were non-white, and the median SSc disease duration (defined as the duration between first non-Raynaud SSc manifestation and date of initial presentation) was 3.9 years. The characteristics of the validation cohort were similar to those of the discovery cohort except for a lower proportion of non-whites, higher prevalence of PAH, lower prevalence of ILD, larger left ventricular (LV) end-diastolic volume, and slightly smaller right ventricular (RV) end-diastolic area in the validation cohort. In both discovery and validation cohorts, the proportion of patients with lcSSc was ~60% with the remaining ~40% of patients with dcSSc.

**Table 1.** Demographic, Clinical, Laboratory, and Echocardiographic Characteristics: Discovery (Northwestern) vs. Validation (Utah) Cohorts

| Characteristic | Training SSc cohort<br>(N=316) | Validation SSc cohort<br>(N=67) | P-value |
| --- | --- | --- | --- |
| Age | 51±12 | 54±13 | 0.21 |
| Female sex, n(%) | 265(84) | 55(82) | 0.72 |
| Race, n(%) |  |  | 0.02 |
| • White | 224(71) | 57(87) |  |
| • Non-white | 92(29) | 10(15) |  |
| SSc disease duration, years* | 3.9(1.4-10.0) | 5(2-10) | 0.03 |
| mRSS (skin score)* | 7(3-18) | 5(4-13) | 0.18 |
| SSc subtype, n(%) |  |  | 0.56 |
| • Limited cutaneous SSc | 186(59) | 42(63) |  |
| • Diffuse cutaneous SSc | 130(41) | 25(37) |  |
| Autoantibodies |  |  |  |
| • Anticentromere antibody, n(%) [n=338] | 66(24) | 16(24) | 0.94 |
| • Anti Scl-70 antibody, n(%) [n=377] | 84(27) | 11(16) | 0.07 |
| Pulmonary arterial hypertension, n(%) | 28(9) | 19(28) | <0.0001 |
| Interstitial lung disease, n(%) | 161(51) | 25(37) | 0.04 |
| Echocardiography data |  |  |  |
| • LV end-diastolic volume, ml | 73±20 | 83±22 | 0.0004 |
| • LV end-systolic volume, ml | 29±13 | 32±11 | 0.08 |
| • LV ejection fraction, % | 61±6 | 62±5 | 0.47 |
| • LV mass, g | 141±45 | 132±44 | 0.18 |
| • E/A ratio | 1.22±0.41 | 1.28±0.41 | 0.29 |
| • e' velocity, cm/s | 9.3±2.8 | 8.7±2.5 | 0.14 |
| • E/e' ratio | 10.1±3.6 | 11.0±5.1 | 0.10 |
| • LA volume, ml | 48±14 | 50±18 | 0.15 |
| • RV end diastolic area, cm<sup>2</sup> | 24±6 | 22±6 | 0.004 |
| • RV wall thickness, mm | 4.6±0.6 | 4.6±0.4 | 0.56 |
| • PA systolic pressure, mmHg | 35±14 | 32±12 | 0.24 |
| • TAPSE, cm | 2.1±0.5 | 2.1±0.3 | 0.47 |

Continuous variables displayed as mean ± standard deviation unless otherwise specified.
SSc = systemic sclerosis; MRSS = modified Rodnan skin score; LV = left ventricular; E = early mitral inflow; A = late (atrial) mitral inflow; e' velocity = early tissue Doppler velocity; LA = left atrial; RV = right ventricular; TAPSE = tricuspid annular plane systolic excursion
\*Values represent median (25<sup>th</sup>-75<sup>th</sup> percentile)

After imputation of missing echocardiographic data (0.36% of the total echocardiographic data), PCA applied to 56 echocardiographic variables identified 6 new feature vectors that collectively accounted for 64% of the phenotypic variance within the echocardiography dataset. Forced vital capacity, ratio of forced expiratory volume in 1 second to forced vital capacity, and the diffusing capacity of carbon monoxide were identified as the 3 PFT variables that were most informative after filtering out highly correlated (R>0.8) variables. Four distinct clusters (pheno-group #1: n=72/316 [23%]; pheno-group #2: n=117/316 [37%]; pheno-group #3: n=58/316 [18%]; and pheno-group #4: n=69/316 [22%]) were identified using model-based clustering of a combination of echocardiophic, PFT, and clinical variables. Internal validation analyses demonstrated that repeating the model-based clustering expectation-maximization algorithm 50 times resulted in the identification of 4 as the most commonly selected optimal number of pheno-groups (36/50 [72%]).

**Table 2** displays the demographic, clinical, PFT, and laboratory characteristics stratified by pheno-group, and **Table 3** display the echocardiographic characteristics for each of the 4 pheno-groups. These results demonstrate that the 4 pheno-groups differ widely in clinical and echocardiographic parameters: each pheno-group represents a unique SSc subtype with characteristic clinical and echocardiographic findings.

**Table 2.**
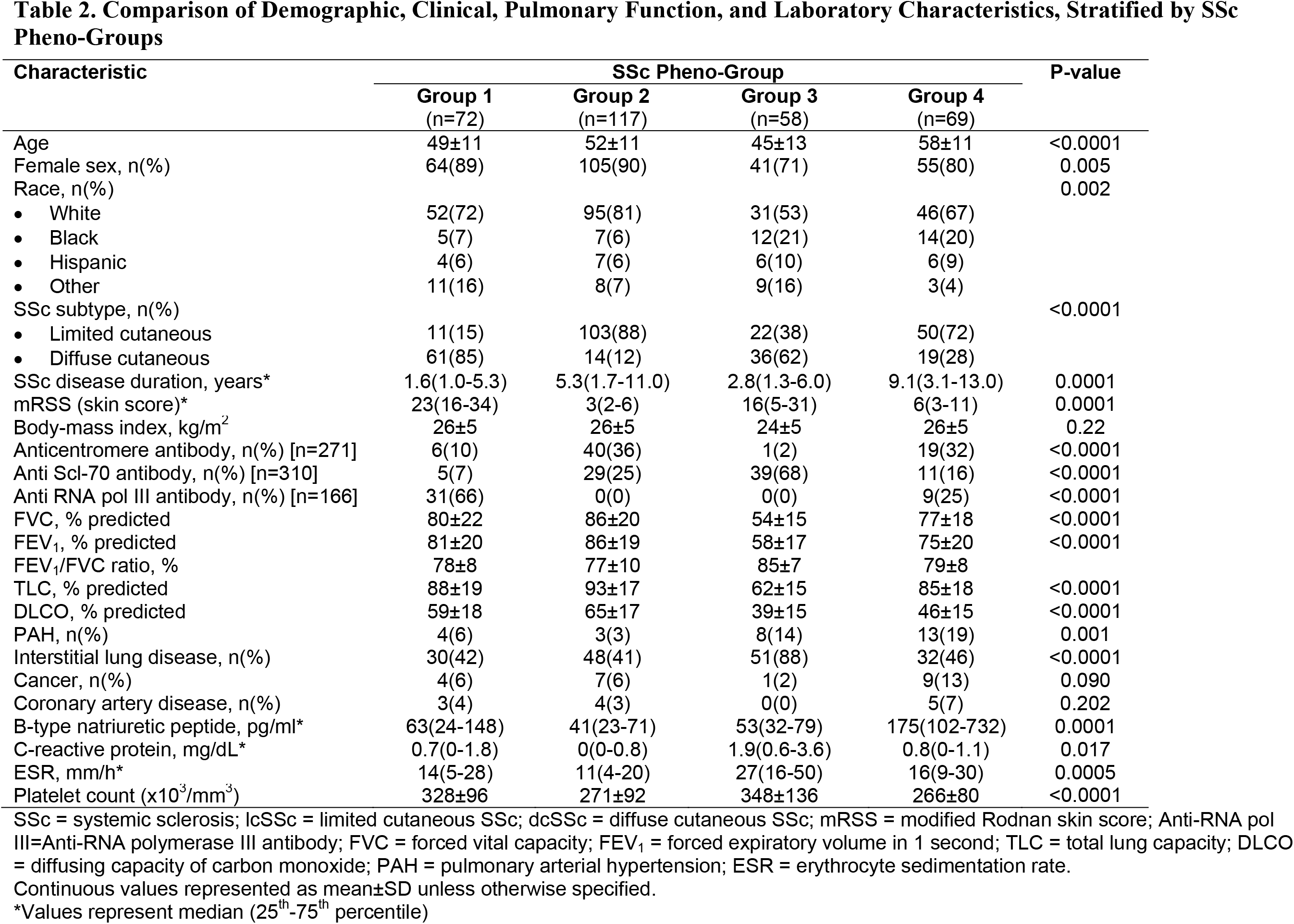
Comparison of Demographic, Clinical, Pulmonary Function, and Laboratory Characteristics, Stratified by SSc Pheno-Groups

**Table 3.** Comparison of Echocardiographic Characteristics, Stratified by SSc Pheno-Groups

| Echocardiographic parameter | SSc Pheno-Group |  |  |  | P-value |
| --- | --- | --- | --- | --- | --- |
|  | Group 1<br>(n=72) | Group 2<br>(n=117) | Group 3<br>(n=58) | Group 4<br>(n=69) |  |
| Systolic blood pressure, mmHg* | 117±22 | 119±15 | 113±17 | 122±22 | 0.06 |
| Diastolic blood pressure, mmHg* | 74±15 | 72±9 | 70±9 | 72±13 | 0.26 |
| Heart rate, bpm* | 77±12 | 70±9 | 82±14 | 76±13 | <0.0001 |
| LV end-diastolic volume, ml | 73±16 | 72±15 | 74±20 | 75±30 | 0.70 |
| LV end-systolic volume, ml | 28±9 | 27±7 | 28±8 | 34±24 | 0.008 |
| LV ejection fraction, % | 62±5 | 62±3 | 62±5 | 58±10 | 0.0003 |
| LV mass, g | 138±34 | 132±34 | 138±44 | 163±65 | 0.005 |
| E wave, cm/s | 91±17 | 87±19 | 84±17 | 86±24 | 0.27 |
| A wave, cm/s | 75±18 | 72±17 | 76±19 | 86±28 | 0.0003 |
| E/A ratio | 1.3±0.38 | 1.3±0.38 | 1.2±0.36 | 1.1±0.51 | 0.036 |
| e' velocity, cm/s | 10.1±2.6 | 9.9±2.4 | 9.3±2.6 | 7.5±2.9 | <0.0001 |
| E/e' | 9.6±3.1 | 9.2±2.5 | 9.5±2.6 | 12.8±5.0 | <0.0001 |
| LA volume, ml | 48±12 | 47±11 | 45±11 | 50±19 | 0.12 |
| RV end diastolic area, cm<sup>2</sup> | 23±5 | 23±5 | 25±6 | 27±9 | 0.001 |
| RV wall thickness, mm | 4.5±0.3 | 4.5±0.4 | 4.7±0.5 | 5.1±0.8 | <0.0001 |
| RA area, cm<sup>2</sup> | 17±4 | 17±4 | 17±4 | 20±8 | 0.005 |
| PA systolic pressure, mmHg | 30±6 | 31±7 | 35±12 | 46±21 | <0.0001 |
| TAPSE, cm | 2.4±0.5 | 2.3±0.4 | 2.0±0.4 | 1.9±0.6 | <0.0001 |
| RV fractional area change, % | 46±4 | 47±4 | 43±7 | 40±9 | <0.0001 |

SSc = systemic sclerosis; lcSSc = limited cutaneous SSc; dcSSc = diffuse cutaneous SSc; LV = left ventricular; E = early mitral inflow; A = late (atrial) mitral inflow; e' = early diastolic tissue Doppler velocity; LA = left atrial; RV = right ventricular; PA = pulmonary artery; TAPSE = tricuspid annular plane systolic excursion.
Values represented as mean ± SD.
\*Vital signs were obtained at the time of echocardiography.

### SSc cardiac pheno-group #1: Marked skin involvement with minimal cardiac involvement phenotype

Patients in pheno-group #1 were more likely to have dcSSc with the highest mRSS scores, indicating the most severe skin involvement. These patients also had the shortest SSc disease duration. Patients in pheno-group #1 also had the highest prevalence of anti-RNA polymerase III antibodies, which are known to be associated with significant skin involvement in SSc. As shown in **Table 2**, for virtually every echocardiographic parameter, pheno-group #1 patients had the most normal values indicating minimal cardiac involvement.

### SSc cardiac pheno-group #2: Limited skin involvement, long SSc disease duration, and mild cardiac involvement phenotype

Patients in pheno-group #2 were more likely to have limited skin involvement (predominantly lcSSc with low mRSS skin scores). In addition, pheno-group #2 had the prolonged SSc disease duration. Both LV and RV involvement were quite mild in this group, with no major echocardiographic abnormalities.

### SSc cardiac pheno-group #3: Inflammatory, interstitial lung disease-right ventricular dysfunction phenotype

Patients in pheno-group #3 had the highest percentage of males and African Americans, most often had diffuse skin involvement (dcSSc), and had a much higher prevalence of interstitial lung disease (88%) than the other 3 pheno-groups (41–46%). These patients also had higher heart rates and significant RV remodeling and dysfunction with relatively preserved LV structure and function. In addition, circulating markers of inflammation (erythrocyte sedimentation rate and C-reactive protein) were highest in pheno-group #3.

### SSc cardiac pheno-group #4: Older, diffuse cutaneous involvement with biventricular dysfunction phenotype

Pheno-group #4 patients were the oldest, were more frequently African American, had limited skin involvement (lcSSc), and had the longest disease duration (median 9.1 years, which was nearly 4 years longer than any of the other pheno-groups). These patients had the greatest cardiac involvement with the highest BNP values, worst LV systolic function (higher LV end-systolic volume, lowest LV ejection fraction), and the most LV hypertrophy (highest LV mass). LV diastolic dysfunction was also most prominent in pheno-group #4 (worst myocardial relaxation [lowest e’ velocity] and highest LV filling pressures [highest E/e’ ratio]). In addition, these patients had significant RV dysfunction with the highest right atrial and pulmonary artery systolic pressures among the 4 pheno-groups (**Table 3**).

### The SSc cardiac pheno-group classification is independently associated with mortality

Mortality was lowest in pheno-groups #1 and #2 and highest in pheno-groups #3 and #4 (**Figure 2** and **Table 4**). We fit a series of nested Cox proportional hazards models to evaluate whether the pheno-groups added prognostic value to established risk factors (**Table 4**). After adjustment for age, sex, race, SSc type, SSc disease duration, PAH, and ILD, pheno-group #3 (hazard ratio [HR] 7.8, 95% confidence interval [CI] 1.5–33.0) and pheno-group #4 (HR 10.5, 95% CI 2.1–52.7) remained associated with mortality (P<0.05 for both pheno-groups). These associations persisted after adjusting for the top 6 echocardiographic principal components suggesting that pheno-group classification provides additional prognostic information beyond that imparted by echocardiographic data alone (**Table 4**).

**Figure 2.**
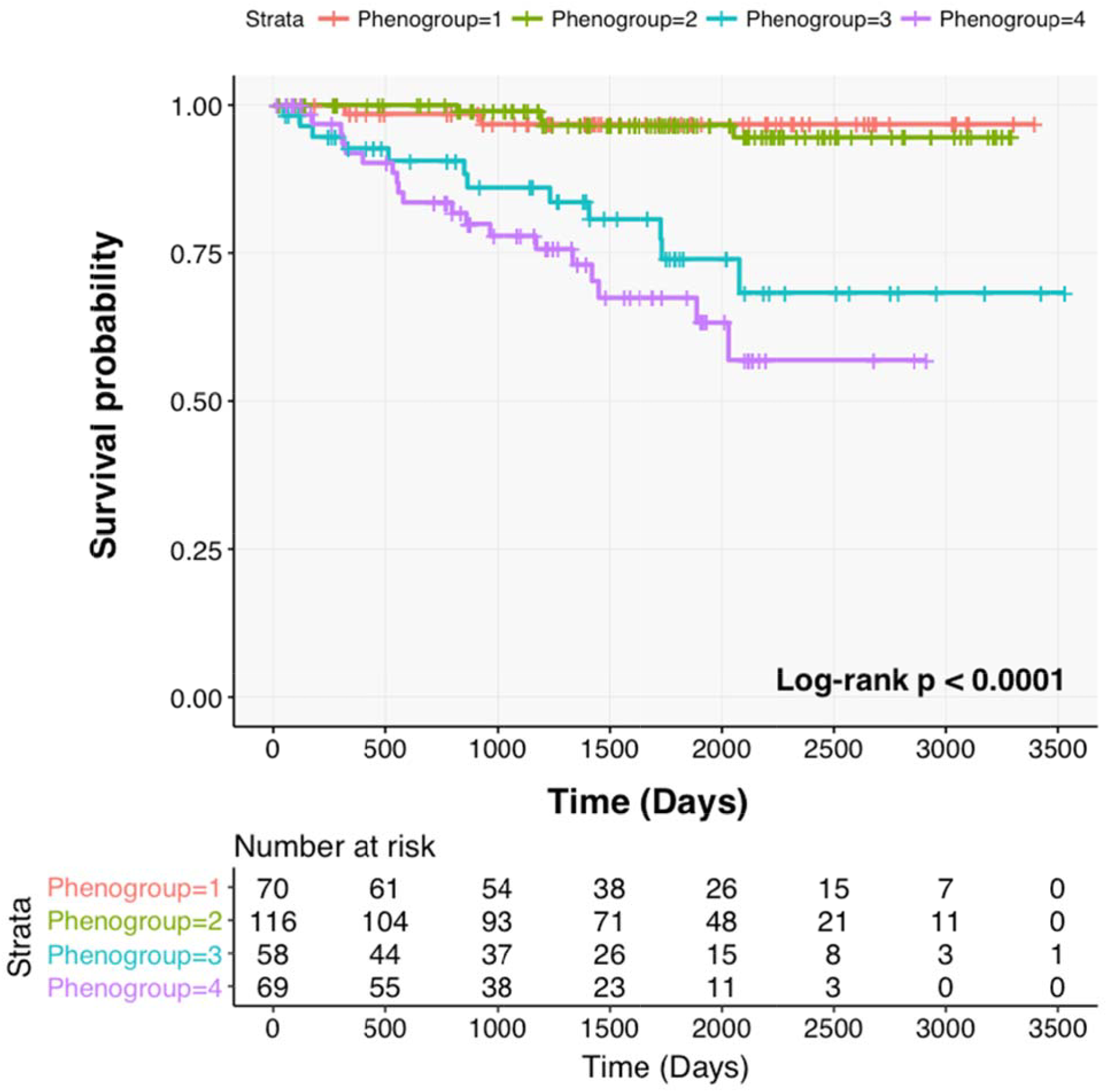
Kaplan-Meier Survival Curves Stratified by SSc Pheno-Group in the Discovery Cohort. dcSSc = diffuse cutaneous systemic sclerosis; lcSSc = limited cutaneous systemic sclerosis

**Table 4.** Association of Pheno-groups with Mortality on Cox Proportional Hazards Analysis

|  | SSc Pheno-Group |  |  |  | P-value for trend | LR test p-value<sup>†</sup> |
| --- | --- | --- | --- | --- | --- | --- |
|  | Group 1<br>(n=72) | Group 2<br>(n=117) | Group 3<br>(n=58) | Group 4<br>(n=69) |  |  |
| Mortality, n(%) | 2(3) | 4(3) | 12(21) | 19(28) | <0.001 | — |
| Unadjusted, HR (95% CI) | 1.0 | 1.2 (0.2-6.4) | 8.9 (2.0-39.6)** | 13.4 (3.1-57.7)*** | <0.001 | — |
| Model 1, HR (95% CI) | 1.0 | 1.4 (0.2-8.5) | 8.8 (1.9-40.3)** | 11.4 (2.4-54.3)** | <0.001 | <0.0001 |
| • Adjusted for age, sex, SSc type, and SSc disease duration |  |  |  |  |  |  |
| Model 2, HR (95% CI) | 1.0 | 1.6 (0.3-9.9) | 7.8 (1.5-33.0)* | 10.5 (2.1-52.7)** | <0.001 | 0.0002 |
| • Adjusted for Model 1 covariates plus race, PAH, and ILD |  |  |  |  |  |  |
| Model 3, HR (95% CI) | 1.0 | 2.6 (0.4-16.7) | 7.3 (1.5-36.6)* | 5.8 (1.0-32.6)* | 0.022 | 0.038 |
| • Adjusted for Model 2 covariates plus the top 6 echocardiographic principal components |  |  |  |  |  |  |

\*P<0.05; \*\*P<0.01; \*\*\*P<0.001
†Likelihood ratio (LR) test p-value for the comparison of the Cox regression model with vs. without the pheno-group classification
HR = hazard ratio; CI = confidence interval; SSc = systemic sclerosis; PAH = pulmonary arterial hypertension; ILD = interstitial lung disease

Although our unsupervised model-based clustering analysis was performed blinded to outcomes, we found that all models that included the pheno-group classification improved prognostication over conventional Cox regression models (by likelihood ratio test), even when they included echocardiographic principal components as covariates. However, the improvement decreased as the model included more and more features that had been used to define patient clusters (pheno-groups), as shown in **Table 4**.

In the independent external validation cohort of 67 patients (University of Utah), we assigned individuals into the 4 pheno-groups as described above. Given the smaller sample size, we compared mortality between the lowest-risk groups (pheno-group #1 and #2) to the highest-risk groups (pheno-groups #3 and #4). Although the Kaplan-Meier curves of the low-risk vs. high-risk pheno-groups were similar to the discovery cohort and showed clear separation over time, the log-rank p-value was not significant (p=0.17) in the setting of low event rates (**Supplementary Figure S1**). However, in the validation cohort, inclusion of the pheno-group classification into the multivariable model (age, sex, race, SSc type, SSc disease duration, PAH, and ILD) again resulted in improvement over a model of conventional risk factors (p=0.008, likelihood ratio test).

As shown in the **Supplementary Material** section, the random survival forests (supervised learning) model identified several known features that are associated with poor outcome in SSc (including indicators of ILD and PAH). In addition to parameters associated with ILD and PAH, pheno-group classification was among the most important predictors of survival, even when including all of the echocardiographic variables in analysis (as shown in the variable importance plot of the random survival forests model [**Supplementary Figure S2**]).

### WGCNA of the skin gene expression in SSc identifies pathophysiologically relevant gene modules

Skin biopsy microarray data from 68 SSc patients, enabled integration of cardiac phenomics with skin gene expression, thereby providing a molecular probe into differences in SSc cardiac disease pathogenesis among the 4 pheno-groups (**Figure 1**). After filtering of gene expression transcripts based on low coefficient of variation in skin biopsies in control subjects and high coefficient of variation in SSc subjects (**Supplementary Figure S3**), WGCNA identified 9 co-expression gene modules within SSc skin (**Supplementary Figure S4**). As shown in **Figure 3A**, only modules 2 and 9 were significantly different among the 4 SSc pheno-groups; whereas modules 2, 6, 7, and 9 were discriminative of affected vs. unaffected skin and differed between lcSSc and dcSSc (**Supplementary Figure S5**). To characterize the molecular pathways underlying these modules, we performed GO association analysis (**Figure 3B**). Module 2, which strongly discriminated clinically involved from non-clinically involved skin, primarily reflects fibrosis, with genes involved in collagen fibril organization and fibroblast growth factor receptor signaling. Other enriched terms included those involving angiogenesis and plasminogen activation, reflecting the vascular contribution to SSc pathogenesis. Module 6 captures stimulation of the innate and adaptive immune response, including T lymphocyte and macrophage activation, Toll-like receptor activity, and response to interferon gamma. Module 9, which is inversely correlated with mRSS, had the enriched term “saliva secretion” and “secretion by tissue”, which we believe reflects eccrine sweat gland architecture (and therefore skin integrity), as demonstrated by inclusion of genes known to be involved in sweat production and secretion, such as the critical sweat gland aquaporin *AQP5*^37^ and the Na-K-Cl cotransporter *SLC12A2*,^38^ as well as the muscarinic cholinergic receptors *CHRM2*^39^ and *CHRM3*.^40^ As is sometimes seen with module annotation, we did not find any enriched GO terms for module 7. However, a manual examination of the genes contained revealed genes involved in keratinocyte differentiation (*FABP5/E-FABP*)^41^ and oxidative stress (*GSTM2*).^42^

**Figure 3.**
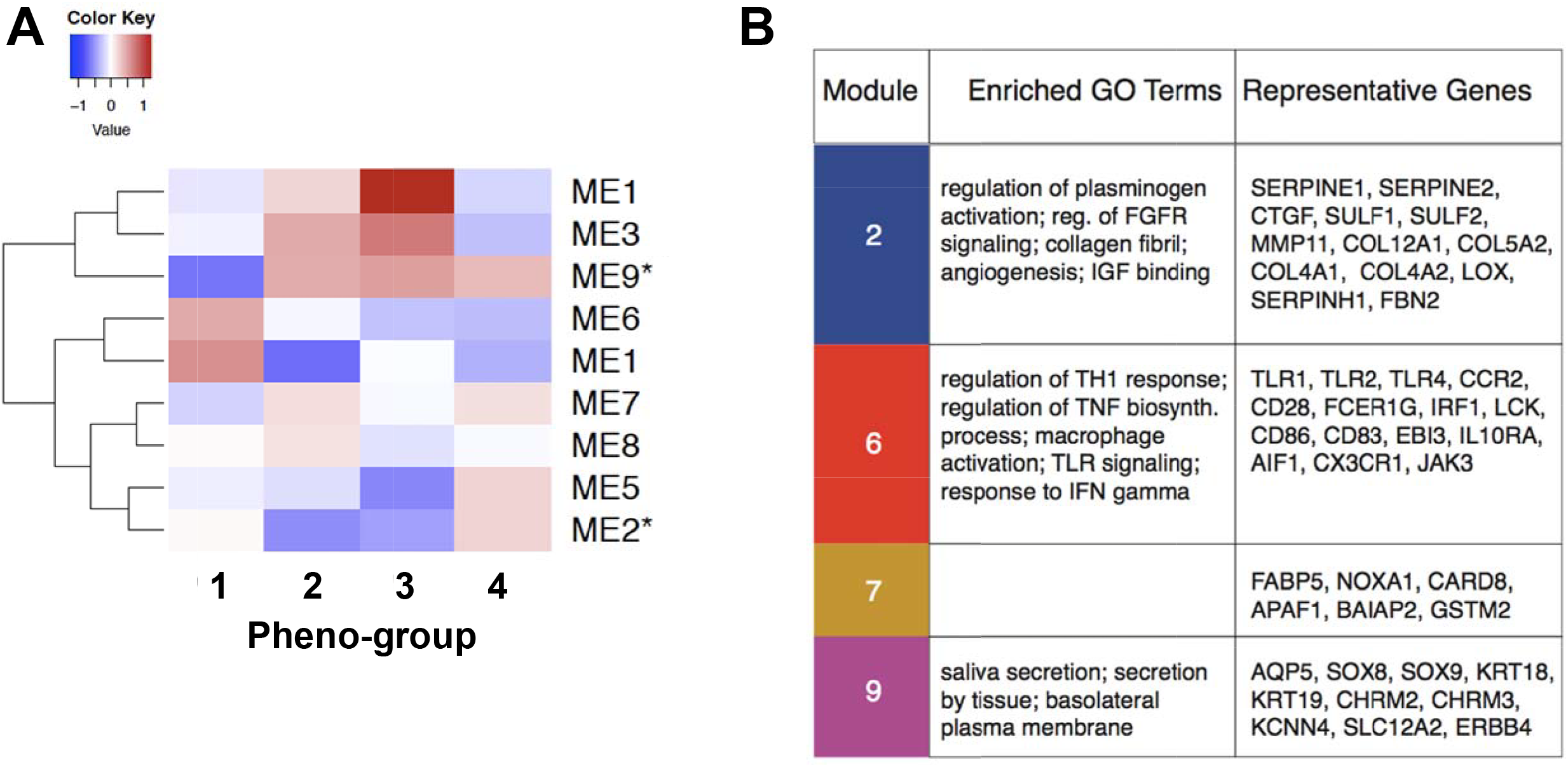
Hierarchical Clustering Heat Map of SSc Pheno-Groups vs. SSc Skin Biopsy Gene Modules. (A) Heat map; (B) representative genes within each of the 4 gene modules that differed among pheno-groups. *p<0.01

Principal component analysis applied to the genes that comprised each module was used to derive a quantitative measure of module activity for each patient. In turn, the correlation between this module score and SSc disease characteristics was computed. Such analysis reinforced the importance of gene modules 2, 6, 7, and 9. We found that modules 2, 6, and 7 in dcSSc and only module 2 in lcSSc readily discriminated clinically affected and non-clinically affected skin in SSc (**Supplementary Figure S5**). Furthermore, module 2, 7 and 9 distinguished dcSSc from lcSSc patients. Modules 2 and 9 were also associated with clinical characteristics of SSc patients (**Figure 4A**), including serum autoantibody profiles (anti-RNA polymerase III, anti-Scl70, and anticentromere antibodies) and mRSS.

**Figure 4.**
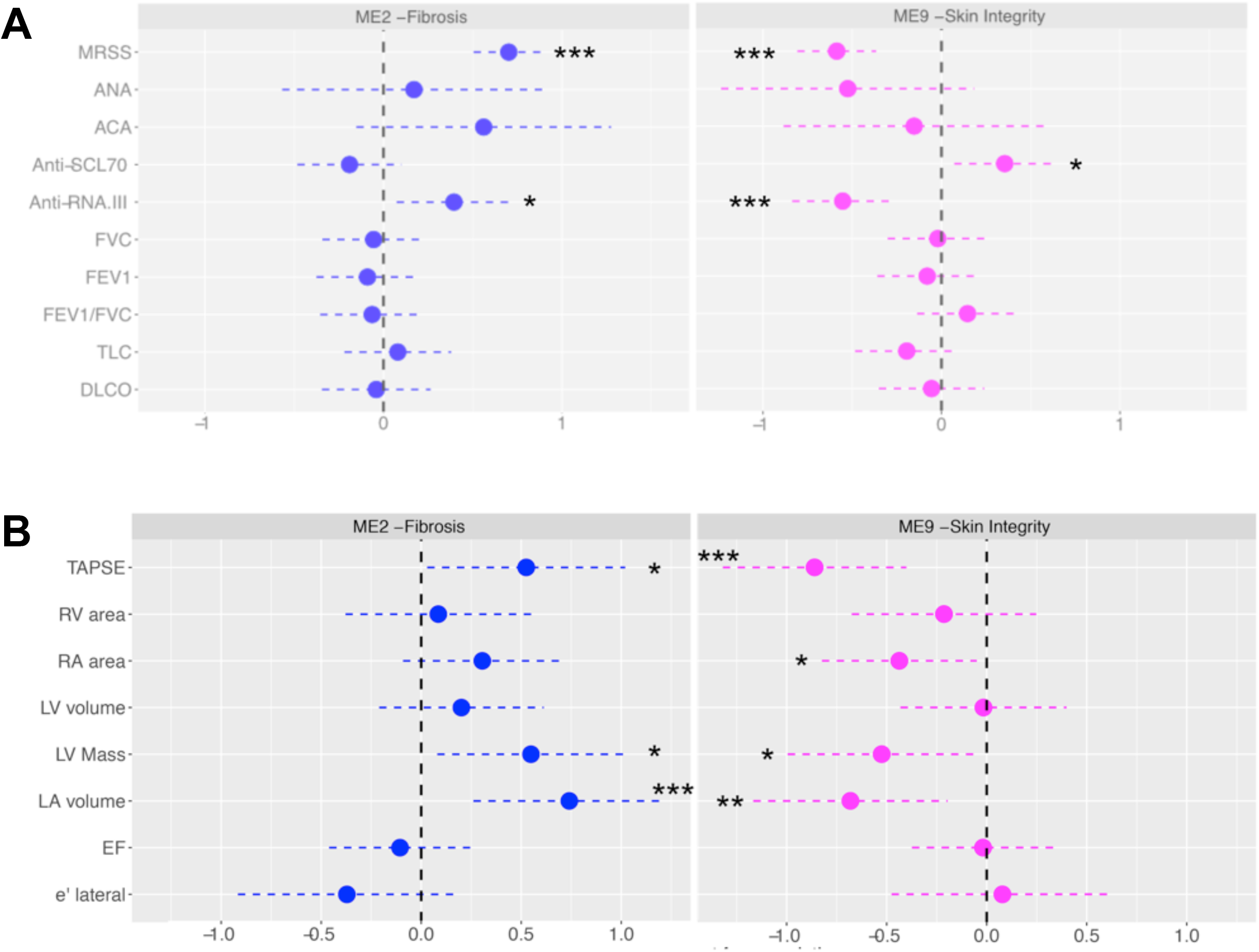
Association of SSc Skin Biopsy Gene Modules with SSc Clinical and Echocardiographic Traits. (A) SSc clinical traits; (B) SSc echocardiography traits. mRSS = modified Rodnan skin score; ACA = anticentromere antibody; ANA = anti-nuclear antibody; anti-RNA.III = anti-RNA polymerase III antibody; anti-SCL70 = anti-Scl70 antibody; DLCO = diffusing capacity of carbon monoxide; FEV1 = forced expiratory volume in 1 second; FVC = forced vital capacity; TLC = total lung capacity; e’ velocity = early diastolic tissue velocity; EF = ejection fraction; LA = left atrial; LV = left ventricular; RA = right atrial; RV = right ventricular; TAPSE = tricuspid annular plane systolic excursion. *p<0.05; **p<0.01; ***p<0.005.

### SSc skin gene expression modules are associated with specific echocardiographic and clinical phenotypes

Next, we investigated whether the various skin gene expression modules were associated with pheno-groups and individual echocardiographic parameters. Pheno-group #1, which had the most extensive skin involvement based on mRSS, had the lowest activity for gene module #9 (likely related to poor skin integrity), whereas pheno-group #4, which had the most extensive cardiac involvement, had the highest activity of gene module #2 (likely related to fibrosis) (**Figure 3A**). We next examined the association of these same modules (#2 and #9) with a subset of echocardiographic traits most representative of aspects of ventricular structure and function (both systolic and diastolic) and atrial size (**Figure 4B**). In keeping with the pheno-group clustering, we found module 2 to be associated with larger LA size and/or greater LV mass (as seen in pheno-group #4), whereas module 9 was associated with smaller right atrial size, LV mass, and LA volume.

## DISCUSSION

Using a machine learning approach to analyze the echocardiographic phenome and skin transcriptome in a cohort of 316 SSc patients, we were able to resolve the heterogeneity of SSc-associated cardiac involvement by identifying 4 novel SSc pheno-groups that differed in clinical characteristics, cardiac structure/function, survival, and molecular profile in the skin. Heterogeneous cardiac abnormalities have been noted in SSc patients for decades;^3^ however, no classification system has been developed to categorize SSc cardiac involvement. Our results lay an important foundation for future clinical studies in SSc cardiac disease that could be informed by a rationale classification strategy for subject stratification. The large size of our SSc cohort, along with densely phenotyped echocardiograms, enabled us to take an unbiased approach towards SSc cardiac disease classification. We identified 4 echo-groups with distinct profiles of cardiac structure and function. Although risk prediction was not the primary objective of our work, pheno-groups #3 and #4 demonstrated higher adjusted all-cause mortality rates that supports the clinical utility of our novel approach. Our study exemplifies the power of machine learning to resolve disease heterogeneity and provide biological insight into identified disease subtypes, an essential step towards precision medicine.

### SSc cardiac pheno-groups provide insight into novel SSc subclasses

Pheno-group #3 exemplifies the utility of our machine learning-based phenomapping framework of SSc cardiac disease. Although pulmonary artery systolic pressure was only slightly higher in this group compared to pheno-groups #1 and #2, these patients had significant RV dysfunction based on reduced RV fractional area change and tricuspid annular plane systolic excursion.

In terms of RV dysfunction etiology, it is unclear to what extent coexisting ILD is a driver or simply a marker of coexisting severe cardiac and lung fibrosis. One of the main values of taking a machine learning approach to comprehensive data that include continuous and dichotomous variables is that a simple binary classification of “ILD or not” would have failed to identify this pheno-group, given the relatively high frequency of ILD (41–46%) in the other 3 pheno-groups. Instead, pheno-groups represent complex recurring patterns of features, readily identifiable by an algorithm, but not easily discovered even by experienced clinicians.

Integration of skin gene expression and echocardiographic data revealed a surprising similarity between pheno-groups #2 and #3 in terms of low expression of genes related to fibrosis (gene module #2), as opposed to pheno-group #4 which had increased activity of this fibrosis-related gene module (**Figure 3A**). Thus, while both pheno-groups #3 and #4 had significant RV dysfunction, widespread fibrosis may be driving the cardiac pathology in pheno-group #4 whereas alternative molecular mechanisms may be driving the RV pathology in pheno-group #3 (or there may be de-coupling of skin and cardiac disease molecular mechanisms in these patients). In addition, it should be noted that although pheno-group #4 was most associated with fibrosis-related gene expression in the skin, these patients were clinically categorized as limited cutaneous (lcSSc) and had low mRSS skin scores, underscoring the ability of machine learning to provide novel insight into SSc cardiac disease subtypes.

### Clinical application of SSc cardiac phenomapping

There are 3 primary potential paths towards clinical application of our work. The first path highlights the need and opportunity to identify cardiac pheno-groups that may be at higher risk for adverse outcomes. Both pheno-groups #3 and #4 had very high hazard ratios for death compared to pheno-group #1. Thus, these high-risk pheno-groups may benefit from more frequent evaluation and targeting with novel therapeutics in clinical trials to try to limit cardiac and extra-cardiac disease progression.

The second application represents a potential to refine clinical trial design using our patient pheno-groups. Echocardiograms are routinely performed for SSc patients and available data could be used to subclassify patients into homogenous groups prior to randomization. Clinical trials are unlikely to be successful if underlying pathophysiology differs among patients. Although our pheno-groups do not yet point to an obvious treatable molecular mechanism, they nonetheless could shape recruitment to ensure trials are powered adequately for different patient classes, with further research (e.g., circulating biomarkers, cardiac magnetic resonance imaging, endomyocardial biopsy) that could inform disease mechanisms.

The third path is further exploration of the skin gene expression profiles as a window into potential treatments that may ameliorate both skin and cardiac manifestations of SSc in particular pheno-groups of patients. Repurposing of FDA-approved drugs that target pathways that our multi-tissue analyses delineate to be important in skin and heart disease could expedite the clinical trials process and identify novel SSc therapeutics. For example, our analysis of gene expression in SSc skin identified a module that included AQP5/aquaporin 5, a specific membrane channel protein that has recently been shown to be expressed in adipocytes as well as fibroblasts.^43^ This is intriguing as studies in mouse models of dermal fibrosis suggest that adipocyte-myofibroblast transition may underlie myofibroblast proliferation in SSc,^44^ and PPAR-γ agonists, such as the thiazolidinediones, play a role in adipogenesis^45^ and fibrogenesis.^46^

### Strengths and limitations

Strengths of our study include the large sample size (especially given the relative rarity of SSc); the inclusion of an external validation cohort; the availability of detailed quantitative echocardiographic measurements; comprehensive clinical phenotyping; and, in a subset, gene expression from skin biopsies of affected and unaffected skin in SSc patients and from healthy controls. Furthermore, to our knowledge, this is the first study to use machine learning to analyze multi-organomics as a way to improve classification of a complex cardiovascular disease.

Several limitations should also be considered when interpreting our results. Although we had a large sample size, with replication of aspects of our work in a second, external cohort, larger validation cohorts are needed. Another outstanding question is the influence of time on patient classification given differences in disease duration among the 4 pheno-groups. However, disease duration was not necessarily a marker of disease severity in our study; for example, pheno-group #1 had the shortest disease duration but had extensive skin involvement based on mRSS skin score. Nonetheless, repeated measurements from the same patient will be invaluable to define patterns of SSc cardiac disease progression among the 4 pheno-groups. Furthermore, although we typically have close temporal juxtaposition of skin biopsy, echocardiographic data, and clinical parameters, the data can be acquired over several months, which may complicate interpretation in rapidly progressive cases. Along these lines, in order to achieve a clinically actionable integration, it will be important to characterize patterns of variation in skin gene expression profiles with changing cardiac manifestations serially in SSc patients over time. Finally, clustering by phenotype cannot define predominant underlying biological mechanism(s), but it can be the first step to trying to understand how phenotypically distinct patients differ according to abnormal pathogenic pathways and/or treatment response.

### Conclusions

Machine learning of echocardiographic and skin gene expression data in SSc is able to identify clinically relevant subgroups with distinct cardiac phenotypes, survival, and associated molecular pathways in the skin. Pheno-group #1 displays a predominant cutaneous phenotype without significant cardiac involvement; pheno-group #2 has long-standing SSc with limited skin and cardiac involvement. Both of these SSc subsets have a favorable prognosis. Pheno-group #3 (with its high propensity for ILD, elevated biomarkers of inflammation, and significant right heart remodeling and dysfunction in the setting of diffuse skin involvement) and pheno-group #4 (with long SSc disease duration, limited skin involvement, and marked biventricular cardiac involvement) both had a poor prognosis. The findings of our study highlight the utility of multi-organ/multi-omic machine learning analyses for classification of complex cardiovascular diseases, which could lead to a more targeted therapeutic approach in line with precision medicine.

## SUPPLEMENTAL METHODS

### Echocardiography

All SSc study participants in both the discovery and validation cohorts underwent comprehensive 2-dimensional echocardiography with Doppler and tissue Doppler imaging (TDI) using commercially available ultrasound systems with harmonic imaging (Philips iE33 or 7500, Philips Medical Systems, Andover, MA; or Vivid 7, GE Healthcare, General Electric Corp., Waukesha, WI). Cardiac structure and function were quantified as recommended by the American Society of Echocardiography (ASE).^1–3^

LV end-diastolic and end-systolic volumes, and left atrial volume, were measured in the apical 4-and 2-chamber views using the biplane method of discs. LV ejection fraction was calculated as (LV end-diastolic volume – LV end-systolic volume)/LV end-diastolic volume. LV mass index was calculated using the linear method, as outlined in the ASE guidelines.^1^ LV diastolic function was graded according to published criteria by using mitral inflow characteristics and tissue Doppler e’ velocities.^4^ Tissue Doppler e’ and s’ velocities were measured at the septal and lateral aspects of the mitral annulus. Sample volume size and placement were optimized for all pulse-wave Doppler and tissue Doppler measurements. All Doppler and tissue Doppler measurements were averaged over three beats (five beats for patients in atrial fibrillation).

Right heart parameters were measured on echocardiography according to published guidelines.^3^ Specifically, we measured RV basal diameter, RV length, RV end-diastolic area, RV end-systolic area, RV wall thickness, and right atrial area. Tricuspid annular plane systolic excursion (TAPSE) was also calculated. Lastly, pulmonary artery systolic pressure was measured using the peak tricuspid regurgitation (TR) velocity (to estimate peak TR gradient) and adding that to the estimated right atrial pressure, which was based on size and collapsibility of the inferior vena cava.^3^

All echocardiographic measurements were made by a single experienced sonographer (LBN) blinded to all other data by an experienced research sonographer using ProSolv 4.0 echocardiographic analysis software (FujiFilm; Indianapolis, IN) and verified by an experienced investigator with expertise in echocardiography (SJS). For the phenomapping analyses, we normalized echocardiographic variables describing chamber dimensions by regressing log-transformed echocardiographic parameters on log-transformed height and gender and using the residuals as phenotypic features.^5^

Non-invasive, echocardiographic indices of pressure-volume analyses were calculated as follows. For the estimation of LV compliance, we calculated the end-diastolic pressure-volume relationship (EDPVR) using a single-beat method^6^ with the equation: LVEDP = α(LVEDV)^β^, where LVEDP is the LV end-diastolic pressure and LVEDV is the LV end-diastolic volume. The EDPVR represents the non-linear relationship between ventricular pressure and volume at end diastole and can be estimated non-invasively. The parameters α and β are constants that allow point measurements of the non-linear EDPVR as reviewed elsewhere,^7^ and were calculated for each individual based on their LVEDV and LVEDP (estimated by [11.96 + (lateral E/e’ ratio)*0.596]).^8^ These parameters were then used to calculate the LVEDV at an idealized LVEDP of 20 mmHg (EDV_20_) for each patient.

To evaluate LV contractility, we estimated the end-systolic pressure volume relationship (ESPVR) as represented by the slope of ESPVR (end-systolic elastance [E_es_]).^7^ E_es_ is a load-independent measure of LV contractility, and can be estimated using a single-beat method.^9, 10^ The relationship between end-systolic pressure (P_es_) and end-systolic volume (ESV) was defined as: [P_es_ = E_es_(ESV – V_0_)]. Using 0.9*(systolic blood pressure) at the time of echocardiography as an estimate of P_es_, we estimated V_0_, the volume-axis intercept, and then generated the estimated ESV at an idealized P_es_ of 120 mmHg (ESV_120_) for each patient. The effective arterial elastance (E_a_), which is a measure of systemic arterial stiffness, was estimated using the equation: E_a_ = 0.9*SBP/stroke volume.^11^ Stroke volume (SV) was estimated on echocardiography using the equation: SV = (LV outflow tract diameter/2)^2^ × LV outflow tract velocity time integral. V-A coupling was defined as E_a_/E_es_.^7^

### Microarray analysis of skin biopsies

Skin biopsies from the dorsal surface of the forearm and the flank were obtained from SSc subjects and healthy control subjects, and microarray studies were performed as previously described,^12^ and as detailed here.

Total RNA (300–500 ng) was amplified and labeled according to Agilent Low RNA Input Fluorescent Linear Amplification protocols. Each experimental RNA sample was labeled with Cy3-CTP and competitively hybridized against Universal Human Reference RNA (Stratagene, La Jolla, CA) labeled with Cy5-CTP on Agilent 44,000-element Human microarrays. Microarrays were scanned with a GenePix 4000B scanner and acquired images quantified with the GenePix Pro 5.1 software (Axon Instruments, Foster City, CA). Technical artifacts and poor-quality spots were flagged and excluded from further analysis. Data were loaded to the UNC Microarray Database.

Data were analyzed as log2 of the LOWESS-normalized Cy5/Cy3 ratio. A total of 18,696 probes with fluorescent signal at least 1.5 greater than local background and of good quality in at least 80% of arrays were selected for analysis. Each data table was multiplied by −1 to convert the log2 ratios to Cy3/Cy5 ratios.

All patients had samples from both clinically involved (fibrotic/edematous) skin, taken from the dorsal forearm, and clinically uninvolved skin (not overtly affected), taken from the back. Previously, skin biopsy data had been used in SSc to classify patients based upon gene expression.^12^ Our primary interest, however, was in clustering genes into coexpression modules, thereby generating quantitative measures of pathway activity to associate with clinical and cardiac phenotypes.

Interpreting microarray data from human tissue can be challenging, in part because a large number of transcripts carry information about inter-individual variation due to genetic, environmental factors or technical causes that can be independent of disease process.^13^ Inclusion of such transcripts in analyses aimed at identifying new and clinically relevant disease classification schemes can result in failure to discriminate between samples from the same individual that are known to be histologically distinct. For example, in a recent study, skin biopsies from individuals with psoriasis and eczema were subjected to microarray analyses. Transcriptional profiles of eczema and psoriatic plaques clustered by patient rather than by disease process indicating that the most differentially expressed genes in microarray datasets from skin biopsies may not be related to disease.^14^ We have previously found that filtering and excluding highly variable transcripts identified in control samples was an effective solution to this problem.^15^ Thus, our analysis pursued a 2-step process. Step 1 was to identify transcripts with high coefficient of variation across 18 samples from 9 healthy control individuals. Step 2 was to identify the most variable transcripts within paired SSc samples (arm and back) excluding any transcripts whose expression was also highly variable across control samples (**Supplementary Figure S2**).

### Random Survival Forest Analysis

We sought to further demonstrate the clinical importance of the SSc cardiac pheno-group classification by using alternate survival models to supplement our results from Cox regression analysis. We hypothesized that supervised learning methods that permit more complex variable combinations may be more effective at modeling survival than Cox proportional hazards models, which are restricted to a linear combination of predictors and as well as the proportional hazards assumption. This may be especially important in SSc since there are complex combinations of features that underlie high-risk SSc phenotypic groups (cardiac phenotypes, pulmonary phenotypes, serum autoantibody profiles, etc.).

One such supervised learning method, named random survival forests (RSF), is an adaption of the random forests algorithm for right-censored data.^16^ The random forest algorithm is a type of ensemble learning method where multiple independent decision trees are fit to the data, and the final output is an average over all trees. In the case of classification, each tree “casts a vote” for class assignment for any data point. The independence of trees is achieved by limiting each tree to only a subset of possible features and individuals (the latter is sometimes referred to as bagging).^17, 18^ For survival data, RSF can accommodate right-censoring and estimate an ensemble cumulative hazard function. To understand the relative importance of individual features, we also computed a random survival model using 81 features and ranked their contribution using variable importance (vimp).

## SUPPLEMENTAL RESULTS

### Random survival forest modeling demonstrates the importance of SSc cardiac pheno-groups

**Supplementary Figure S2** displays the variable importance (vimp) plot, which was generated using the random survival forests model. The vimp plot shows the relative importance of a large number of features that were used in the RSF model to predict survival. Parameters associated with PAH (e.g., RV wall thickness, indicative of RV hypertrophy; tricuspid regurgitation gradient, indicative of elevated pulmonary artery systolic pressure; right atrial pressure, indicative of right-sided heart failure; and the diagnosis of PAH itself), and ILD (e.g., diffusing capacity of carbon monoxide and forced vital capacity) were some of the most important predictors of survival, findings which are consistent with the fact that PAH and ILD are leading causes of death in systemic sclerosis. However, pheno-group membership is also very high on the vimp plot, which adds further evidence that the phenomapping classification is a very strong prognostic indicator in SSc.

---

**Supplementary Table 1.** Clinical, Echocardiographic, and Pulmonary Function Test Phenotypes (Features) Included in the Phenomapping (Machine Learning) Analyses

| **Phenotypic Domain** | **Phenotypes** |
| --- | --- |
| Clinical | Sex, race (white or non-white), mRSS (ordinal for each of 17 body areas, resulting in an integer score [0-51]), autoantibodies (anti-topoisomerase I [Scl-70] antibody, anti-RNA polymerase III antibody, anti-centromere antibody) |
| Left heart structure | LV end-diastolic volume, LV end-systolic volume, LV end-diastolic dimension, LV end-systolic dimension, septal wall thickness, posterior wall thickness, relative wall thickness, LV mid-chamber diameter (apical 4-chamber view), LV outflow tract diameter, LV mass, left atrial volume |
| LV systolic function | LV ejection fraction, tissue Doppler s' velocity (septal and lateral), fractional shortening, velocity of circumferential fiber shortening |
| LV diastolic function | Mitral inflow characteristics (E velocity, A velocity, E/A ratio, E deceleration time, IVRT [measured on septal and lateral tissue Doppler tracings]), tissue Doppler characteristics (septal e' and lateral e' velocities; septal a' and lateral a' velocities; septal E/e' and lateral E/e' ratios) |
| Right heart structure | RV basal diameter, RV maximal diameter, RV length, RV outflow tract diameter (in the short axis and long axis), RV wall thickness, RV end-diastolic area, RV end-systolic area, RV/LV maximal diameter ratio, right atrial area |
| RV function | RV fractional area change, tricuspid annular plane systolic excursion |
| Hemodynamics | LV outflow tract velocity time integral, RV outflow tract velocity time integral, stroke volume, cardiac output, PA systolic pressure, RA pressure |
| Pressure-volume analysis | Effective arterial elastance, end-systolic elastance, systolic blood pressure/end-systolic volume ratio, end-diastolic elastance, ventricular-arterial coupling, stroke work, stroke work/end-diastolic volume ratio, preload recruitable stroke work, pulse pressure/stroke volume ratio |
| Miscellaneous echocardiography variables | Heart rate at the time of echocardiography, pre-ejection period, ejection time |
| Pulmonary function test | FEV1/FVC ratio, FVC, diffusing capacity of carbon monoxide |

LV = left ventricular; RV = right ventricular; IVRT = isovolumic relaxation time; PA = pulmonary artery; RA = right atrial; FEV1 = forced expiratory volume in 1 second; FVC = forced vital capacity

**Supplementary Figure S1.**
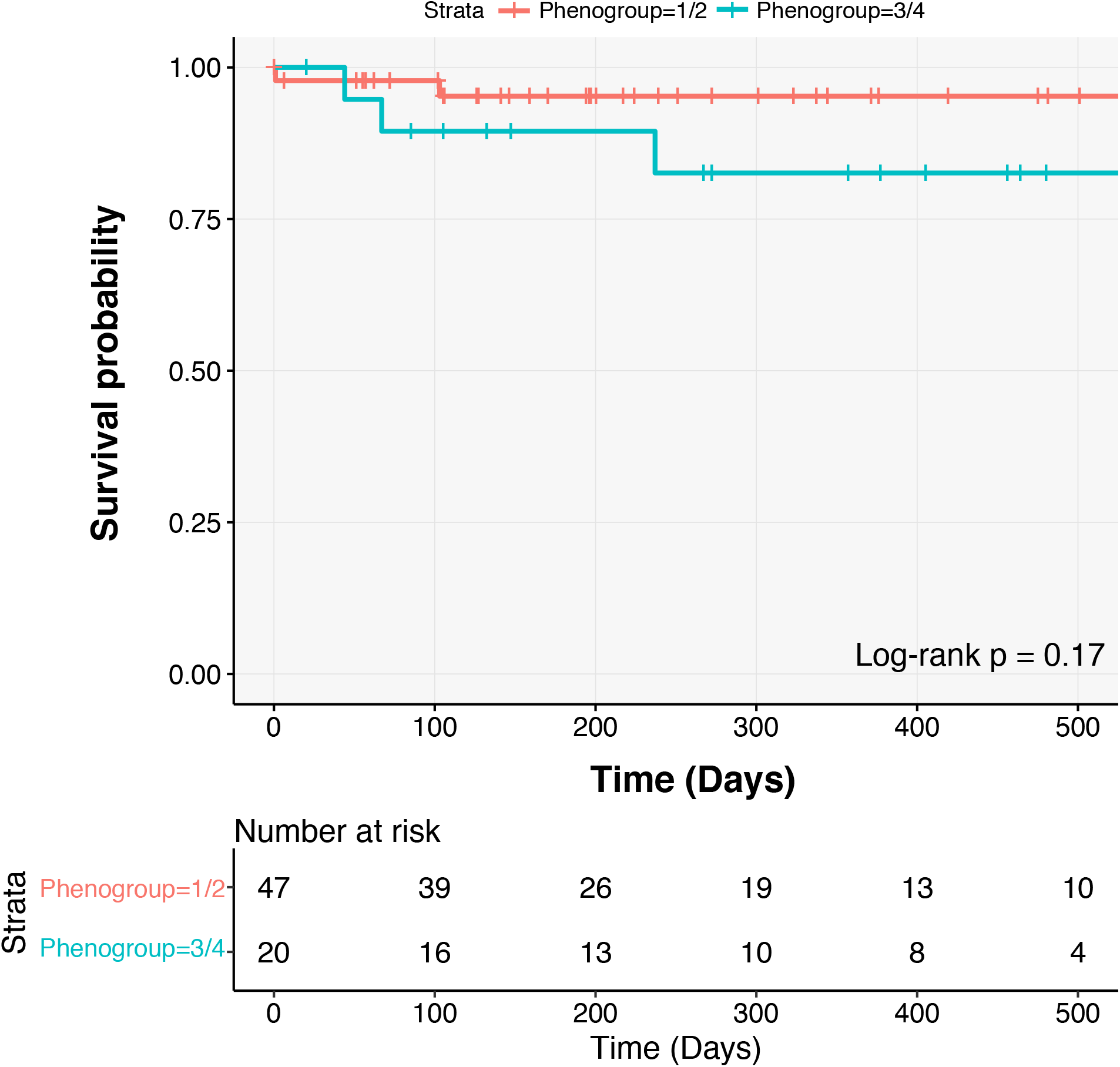
Kaplan-Meier Survival Curves Stratified by Low-Risk and High-Risk SSc Pheno-Groups in the Validation Cohort. In the validation cohort (University of Utah), pheno-groups #1 and #2 were collapsed into one group, and pheno-groups #3 and #4 were collapsed into another group, resulting in 2 groups. The Kaplan-Meier curve shows that the survival trajectories of the “low-risk” and “high-risk” pheno-groups is retained in the validation cohort, although the log-rank P did not achieve statistical significance due to the relatively low number of deaths in the validation cohort. Of the 67 Utah SSc patients, 3/20 (15%) in high-risk pheno-group #3/4 died during follow-up vs. 2/45 (4.2%) in the low-risk pheno-group #1/2 (p=0.15 by chi-squared test; HR 3.3 [95% CI 0.5–19.5], p=0.20 on Cox regression analysis).

**Supplementary Figure S2.**
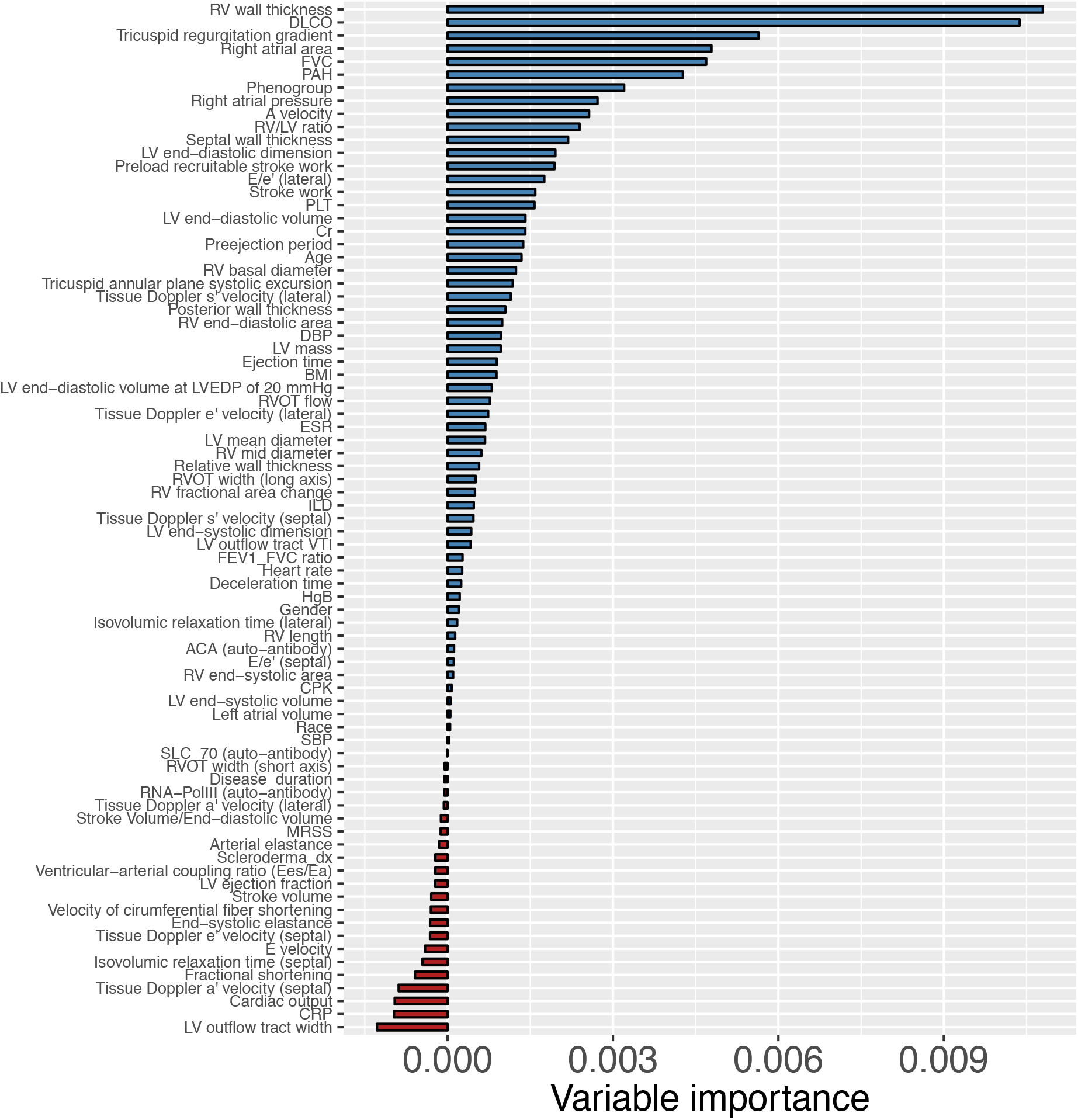
Variable Importance (Vimp) Plot, Based on the Random Survival Forests (RSF) Model. The vimp plot shows the relative importance of a large number of features that were used in the RSF model to predict survival. Parameters associated with pulmonary arterial hypertension (e.g., RV wall thickness, indicative of RV hypertrophy; tricuspid regurgitation gradient, indicative of elevated pulmonary artery systolic pressure; right atrial pressure, indicative of right-sided heart failure; and the diagnosis of pulmonary arterial hypertension itself), and interstitial lung disease (e.g., diffusing capacity of carbon monoxide and forced vital capacity) were some of the most important predictors of survival, which is consistent with the fact that pulmonary arterial hypertension and interstitial lung disease are leading causes of death in systemic sclerosis. However, pheno-group membership is very high on the variable importance plot, along with these other indices, thereby supporting the phenomapping classification as a very strong prognostic indicator in systemic sclerosis.

**Supplementary Figure S3.**
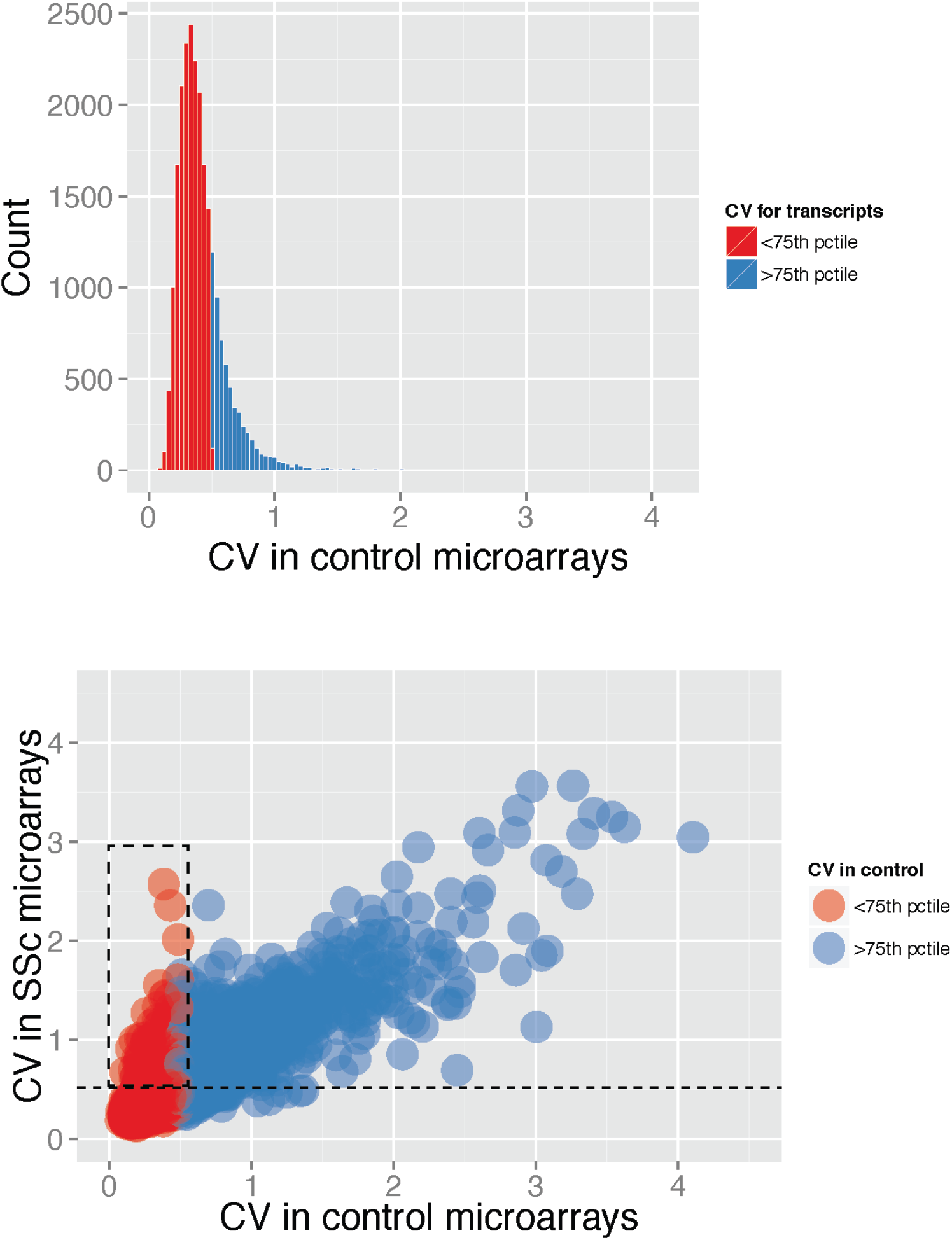
Filtering of Gene Expression Transcripts Based on Low Coefficient of Variation in Skin Biopsies in Control Subjects and High Coefficient of Variation in SSc Subjects. Top panel = identification of transcripts with low (<75^th^ percentile) coefficient of variation in the control skin biopsies; bottom panel = identification of highly variable transcripts in SSc biopsies that have low variability in control skin biopsies.

**Supplementary Figure S4.**
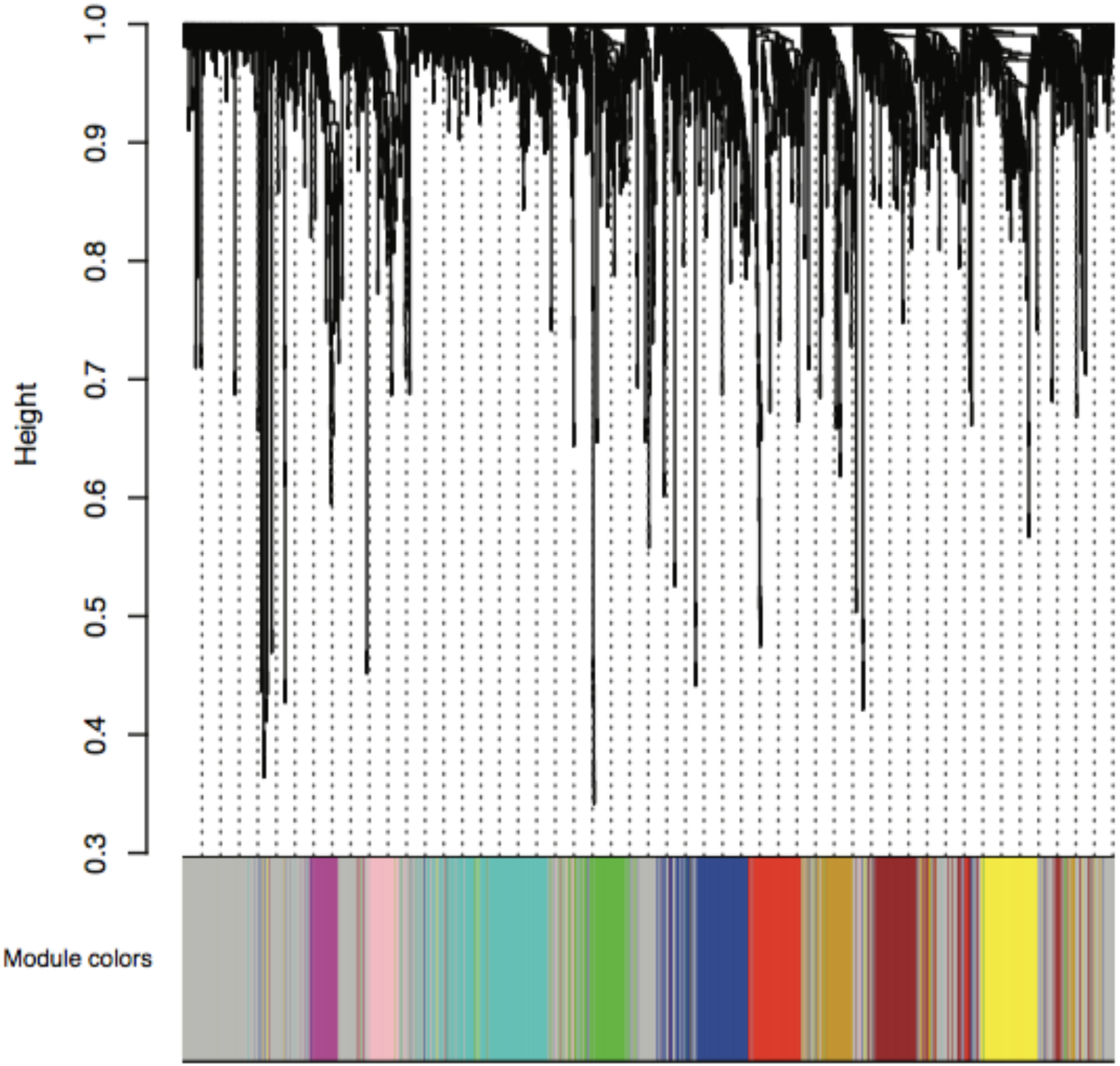
Clustering of SSc Skin Biopsy Genes into Gene Modules Based on Whole Genome Co-Expression Network Analyses

**Supplementary Figure S5.**
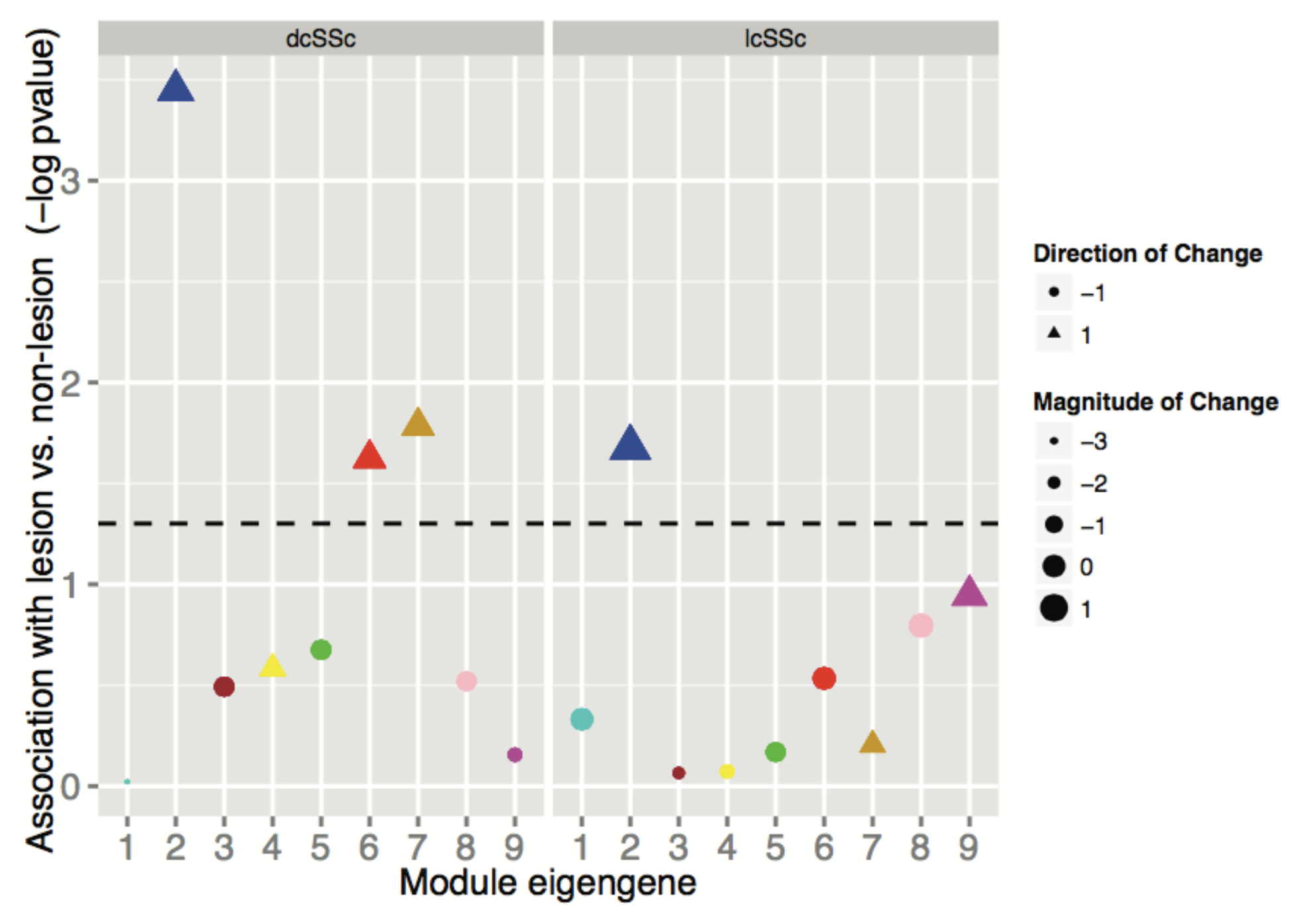
Comparison of the Association of Skin Gene Expression Modules with Lesional vs. Non-Lesional Skin in SSc Patients. In dcSSc, gene modules 2, 6, and 7 were significantly differentially expressed in lesional vs. non-lesional skin in SSc patients. In lcSSc, gene module 2 was significantly differentially expressed in lesional vs. non-lesional skin in SSc patients.

